# Characterization of the zebrafish *gabra1^sa43718/sa43718^* germline loss of function allele confirms a function for Gabra1 in motility and nervous system development

**DOI:** 10.1101/2023.01.27.525860

**Authors:** David Paz, Nayeli G. Reyes-Nava, Briana E. Pinales, Isaiah Perez, Claudia B. Gil, Annalise V. Gonzales, Brian Grajeda, Igor L. Estevao, Cameron C. Ellis, Victoria L. Castro, Anita M. Quintana

**Affiliations:** Department of Biological Sciences, Border Biomedical Research Center, The University of Texas at El Paso, El Paso, Texas, United States of America

**Keywords:** zebrafish, seizure, GABRA1, GABA

## Abstract

Mutation of the *GABRA1* gene is associated with neurodevelopmental defects and epilepsy. *GABRA1* encodes for the α1 subunit of the gamma-aminobutyric acid type A receptor (GABA_A_R), which regulates the fast inhibitory impulses of the nervous system. Multiple model systems have previously been developed to understand the function of *GABRA1* during development, but these models have produced complex and at times incongruent data. Thus, additional model systems are required to validate and substantiate previously published results. We investigated the behavioral swim patterns associated with a nonsense mutation of the zebrafish *gabra1* (*sa43718* allele) gene. The *sa43718* allele causes a decrease in *gabra1* mRNA expression, which is associated with light induced hypermotility, one phenotype associated with seizure like behavior in zebrafish. Mutation of *gabra1* was accompanied by decreased mRNA expression of *gabra2, gabra3, and gabra5,* indicating a reduction in the expression of additional alpha sub-units of the GABA_A_R. Although multiple sub-units were decreased in total expression, larvae continued to respond to pentylenetetrazole (PTZ) indicating that a residual GABA_A_R exists in the *sa43718* allele. Proteomics analysis demonstrated that nonsense mutation of *gabra1* is associated with abnormal expression of proteins that regulate proton transport, ion homeostasis, vesicle transport, and mitochondrial protein complexes. These data support previous studies performed in a zebrafish nonsense allele created by CRISPR/Cas9 and validate that loss of function mutations in the *gabra1* gene result in seizure like phenotypes with abnormal function of inhibitory synapses.

## Introduction

The gamma-aminobutyric acid type A receptor (GABA_A_R) is a multi-subunit ion channel that mediates inhibitory synapses of the nervous system. The GABA_A_R can be composed of unique combinations of any of the following: 6 α subunits, 3 β subunits, 3 γ subunits, and one ε, δ, θ, or π subunits. The most common GABA_A_R in mammals consists of 2 α subunits, 2 β subunits, and γ subunit. Each of the subunits is encoded by an independent gene, each of which are located on different chromosomes. Mutations in many of these subunits have been associated with seizure phenotypes in humans ^1^ and most recently we used whole exome sequencing to identify a putative heterozygous mutation in the *GABRA1* gene (c.875C>T) associated with seizure phenotypes ^2^. *GABRA1* encodes for the α1 subunit of the GABA_A_R and to date, over 30 different variants in the *GABRA1* gene have been reported and associated with neurological disorders and neurodevelopmental defects ^2–11^. Subsequent studies have demonstrated that the c.875C>T mutation is a loss of function allele that reduces single channel open time ^12^.

Several murine models have been developed to investigate the physiological and behavioral functions of *GABRA1* in mammals, however, the phenotypic outcomes are heterogeneous. For instance, deletion of *Gabra1* or the knock-in of the p.Ala332Asp allele cause behavioral phenotypes that range from tremors/absence-like seizures to the development of myoclonic seizures at postnatal stages ^13–16^. These phenotypes have been strain, age, and sex specific, which makes interpretation of results difficult. Additional zebrafish models have been characterized including a germline mutant and a morpholino mediated transient knockdown. These two studies yielded contrasting results whereby hypermotility was observed in a nonsense mutant of *gabra1* and hypomotility was observed after transient knockdown ^2, 17^. Thus, additional validatory models are required to understand the function of *gabra1* during early development .

In 2016, the zebrafish mutation project was completed by the Wellcome Sanger Institute, where over 40,000 mutant alleles, covering 60% of zebrafish protein-coding genes, were generated ^18^. However, only a small number of alleles have been characterized or associated with a specific phenotype. An allele carrying a nonsense mutation in the *gabra1* gene (*sa43718*) was generated through N-ethyl-N-nitrosourea treatment and remains uncharacterized. We hypothesized that a more comprehensive analysis of this allele could discern the disparate phenotypes observed across other zebrafish models (nonsense versus transient knockdown). We therefore performed behavioral and molecular analysis of the sa43718 allele.

Our results indicate that homozygous *gabra1^sa43718/sa43718^* larvae show hyperactive locomotion (seizure-like behavior) upon a light stimulus. Interestingly, despite a significant reduction in *gabra1* mRNA expression, homozygous mutants continued to respond to pentylenetetrazole (PTZ), which binds to and antagonizes the GABA_A_R. Based on this response, we hypothesized that other Gabra alpha sub-units had increased expression and integrated in GABA_A_R resulting in a blunted, but consistent PTZ response. However, we did not detect up-regulation of other alpha sub-units and our analysis revealed reduced expression of *gabra1*, *gabra2*, *gabra3*, and *gabra5*. Consistent with the decrease in multiple alpha sub-units, proteomics analysis identified abnormal expression of proteins that regulate proton transport, sodium ion homeostasis across the plasma membrane, synapse development, and the mitochondrial inner membrane protein complex and respiratory chain, thus suggesting abnormal development of the nervous system inhibitory response.

## Results

### Experimental model and animal husbandry

The *gabra1^sa^*^43718*/+*^ allele (N=10 female fish) was obtained from Sanger Institute through the Zebrafish International Resource Center (ZIRC). Female carriers were outcrossed with wildtype (AB) males to generate independent families of heterozygous male and female carriers. For all experiments, zebrafish larvae were obtained by mating adult heterozygous *gabra1^sa^*^43718*/+*^ fish. Collected zebrafish embryos were maintained in E3 media (5mM NaCl, 0.17mM KCl, 0.33mM CaCl_2_, 0.33mM MgSO4, 0.05% methylene blue, pH 7.4) at 28°C under 14:10 light: dark cycle. All experimental procedures were performed at 5 days post fertilization (DPF). All animals were maintained and used in accordance with the guidelines from the University of Texas at El Paso Institutional Animal Care and Use Committee (Animal Protocol Number 811869-5). Euthanasia and anesthesia were performed according with the American Veterinary Medical Association (AVMA) guidelines for the euthanasia, 2020 edition.

### Genotyping

Genotyping of the *gabra1^sa^*^4371*/sa*4371^ allele was performed by PCR amplification and restriction enzyme digest. Adult fin clips and larval tails were lysed in lysis buffer (30ul) (50mM sodium hydroxide) for 5 minutes at 95°C followed by 10 minutes at 4°C. DNA was then neutralized by addition of 500mM Tris-HCL solution pH 8.0 (6ul) and used directly for PCR. The fragment of interest (237bp) was amplified by standard PCR at an annealing temperature of 60°C (forward: TTGTGACTCAAAGCCACGAG and reverse: TGAGACGAGAACCATCGTCA). PCR amplicon containing the *sa43718* allele region, was digested with XhoI (New England Biolabs) at 37°C according to manufacturer’s guidelines. The mutation present in the *sa43718* inhibits cleavage by XhoI allowing differentiation of wildtype, heterozygous, and homozygous offspring.

### Behavioral analysis and pentylenetetrazole (PTZ) treatment

Behavioral analysis was performed using the ZebraBox (ViewPoint Behavioral Technology, Montreal, Canada). Briefly, embryos were obtained by natural spawning and raised to 5 DPF. Larvae (5 DPF) were individually tracked for swim speed and total distance swam in a 96-well plate. The behavioral analysis consisted of 15 minutes divided into 5-minute intervals of dark/light/dark conditions. All larvae were acclimated to the chamber for 1 hour prior to data acquisition ^2^. Data was collected each minute, for a total of 15 minutes and total distance traveled (mm) and speed (mm/s) was calculated as previously described ^2^. Experiments were performed in biological duplicates using a minimum of N=30 larvae per trial. Treatment with 10mM pentylenetetrazol (PTZ) (Millipore-Sigma) was performed as previously described ^2^. Genotyping of each larvae was performed after behavioral analysis.

### Quantitative real time PCR (qPCR)

Total RNA was isolated from brain homogenates at 5 DPF from wildtype and homozygous larvae using TRIzol reagent (ThermoFisher) and Directzol columns (Zymo) according to manufacturer instructions. RNA was reverse transcribed using the Verso cDNA Synthesis Kit (ThermoFisher) and total RNA (1ug) was normalized across samples. Gene expression was measured in biological triplicates using a pool of at least N=10-12 larvae per biological replicate. For each individual biological replicate, technical replicates were performed as internal controls for the quantitative PCR (qPCR). The Applied Biosystems StepOne Plus machine with Applied Biosystems associated software was used for qPCR analysis. Sybr green (ThermoFisher) based primer pairs were designed for each gene analyzed: *gabra1* (FWD: TTTTGCTCCGAACATTGCCA, REV: CCGAATAGCAGTGGGAAAGC), g*abra2a* (FWD: GATGGCTACGACAACAGGCT, REV: TGTCCATCGCTGTCGGAAAA), *gabra3* (FWD: GCTGAAGTTCGGGAGCTATG, REV: GGAGCTGATGGTCTCTTTGC), *gabra4* (FWD: GACTGCGATGTACCCCACTT, REV: ATCCAGGTCGGAGTCTGTTG), *gabra5* FWD: CATGACAACACCCAACAAGC, REV: CAGGGCCTTTTGTCCATTTA), *gabra6a* (FWD: TCGCGTACCCATCTTTCTTC, REV: CCCTGAGCTTTTCCAGAGTG), *gabrb2* (FWD: CCCGACACCTATTTCCTCAA, REV: TCTCGATCTCCAGTGTGCAG), *gabrg2* (FWD: ACACCCAATAGGATGCTTCG, REV: AGCTGCGCTTCCACTTGTAT). *gapdh* (FWD: GGCAAGCTTACTGGTATGGC, REV: TGAGAGCAATACCAGCACCA) expression was used as a reference gene for 2^ΔΔct^ quantification. Statistical analysis performed using a *t-test*.

### Protein isolation

For protein analysis, total protein was obtained from a pool of whole brain homogenates (n=9) per genotype. Two biological replicates per genotype were collected for a total of 18 brains. Brains were excised and stored for protein isolation in 1X Cell Lysis Buffer (Cell Signaling) with protease inhibitors cocktail (ThermoFisher). DNA was isolated from the tail tissue for genotyping and protein was isolated from brain homogenates using manual homogenization. Protein quantification was performed using Precision Red Protein Assay Reagent (Cytoskeleton) according to manufacturer’s instructions. Samples were then sent to the Biomolecule Analysis and Omics Unit (BAOU) at The University of Texas El Paso for sample processing and proteomic analysis.

### Sample preparation for proteomics

Proteins were purified by trichloroacetic acid (TCA) protein precipitation. Fifty microliters of 100% TCA (Sigma/Millipore - cat# T6399-5G) was added to 200 µL of sample in 1x Cell Lysis Buffer (Safe Seal microcentrifuge tubes, Sorenson BioScience, cat. No. 12030) and incubated at 4°C for 10 minutes. Total protein ranged from 2 – 5 µg per sample. The precipitated proteins were pelleted by centrifugation for five minutes at 14,000 rcf and the supernatant was discarded. The protein pellet was subsequently washed a total of three times with 200 µL LCMS grade acetone by resuspension and pelleting, the acetone supernatant was discarded each time. Residual acetone was evaporated using a heating block at 95°C for 2 minutes. Samples were stored as a pellet at -80°C until processing for proteomic analysis. Stored pellets were resuspended in 8M Urea prior and subjected to enzymatic tryptic digestion using the PreOmics iST sample preparation kit (catalog no. iST 96x P.O.00027). Briefly, denaturation, alkylation, and reduction were performed with 10 µL LYSE buffer per 1 µg of protein and heated at 80 °C for 20 min while being mixed every 5-minutes. Remaining droplets on the cap were given a brief centrifugation (RT; 300 rcf; 10 sec) and samples were sonicated ten times at 30-second on/off intervals for a total of 5 minutes. Fifty microliters of resuspended DIGEST solution were added to the samples for protein digestion. Sample-filled microtubes were gently vortexed, centrifuged and kept for 90 minutes at 37°C in a heating block. Throughout the 90-minute incubation period, samples were gently mixed every 10 minutes. One hundred microliters of STOP solution were added and mixed by pipetting ten times. Samples were then transferred to the cartridge and centrifuged for 1 minute at 3,800 rcf, followed by subsequent 200 µL washes with WASH 1 and WASH 2 solutions according to the manufacturer’s instructions. Peptides were eluted in two cycles of 100 µL of ELUTE solution and dried completely in a vacuum evaporator (Savant; ThermoFisher) for 90 minutes at 45 °C and 100-mTorr and stored at -80°C until LC-MS/MS acquisition.

### Liquid chromatography–tandem mass spectrometry (LC-MS/MS)

Peptides were resuspended in 4% acetonitrile (ACN) with 0.1% formic acid (FA) at a concentration of 1µg/µL. Peptides were separated by a Dionex Ultimate 3000 UHPLC system (ThermoFisher) tandem with a Q-Exactive Plus Hybrid Quadrupole-Orbitrap Mass Spectrometer (ThermoFisher) with Xcalibur software (v. 3.0.63) for data acquisition in positive mode. Peptides were separated on a C18 Acclaim PepMap nanoLC column (75 µm x 50 cm nanoViper, PN 164942, ThermoFisher) equilibrated with 4% solvent B (99.9% acetonitrile, 0.1% formic acid) and 96% solvent A (99.9% H2O, 0.1% formic acid) kept at 55°C throughout the entire acquisition. One microliter of peptides was loaded onto the column for 15-minutes at a flow rate of 0.5 µL/min and eluted with a multi-step gradient. The flow rate was reduced to 0.3µL/min over the course of 15 min, and solvent B was set to 20% over 100-minutes, then increased to 32% and maintained for 20-minutes before increasing to 95% over 1-minute. The column was washed with 95% solvent B at a flow rate of 0.4 µL/min for 4-minutes to remove/clean any remaining peptides. The column was then re-equilibrated with 4% solvent B at 0.5µL/min until a 180-minutes total runtime. Blank injections were added after each biological replicate, using a 60-minute two sawtooth gradient from 4-95% solvent B, and column re-equilibration at 4% solvent B. The mass spectrometer was set to top10 data-dependent acquisition (DDA), with a scan range of 375 to 1500 m/z and a full MS resolution of 70,000; AGC target 3e6. Ions were fragmented with NCE at 27 and collected with an AGC target of 1e5 at 17,500 resolution, 2 m/z isolation window, maximum IT of 60 ms. Charged exclusion ions were unassigned, 1, 6–8, and >8 charges.

### Bioinformatics data analysis

After proteomics analysis, Proteome Discover (PD) 2.5.0.400 (ThermoFisher) was utilized to identify the proteins from each peptide mixture. The database for Danio rerio was downloaded from UniProtKB; http://www.uniprot.org/ on 21 October 2021 with a database 61,623 sequences. A contaminant dataset was run in parallel composed of trypsin autolysis fragments, keratins, standards found in CRAPome repository and in-house contaminants. PD analysis parameters are as follows: false-discovery rate (FDR) of 1%, HCD MS/MS, fully tryptic peptides only, up to 2 missed cleavages, parent-ion mass of 10 ppm (monoisotopic); fragment mass tolerance of 0.6 Da (in Sequest) and 0.02 Da (in PD 2.1.1.21) (monoisotopic). Two-high confidence peptides per protein were applied for identifications. PD dataset was processed through Scaffold Q+S 5.0.1. Scaffold (Proteome Software, Inc., Portland, OR 97219, USA) was used to probabilistically validate protein identifications derived from MS/MS sequencing results using the X!Tandem and Protein Prophet. Data was transferred to Scaffold LFQ (Proteome Software, Portland, Oregon, USA) which was used to validate and statistically compare protein identifications derived from MS/MS search results. A protein threshold of 95%, peptide threshold of 95%, and a minimum number of 2 peptides were used for protein validation. Normalized weighted spectral counts were used when comparing the samples. To ascertain p-values, Fisher’s Exact was run with a control FDR level q * .05 with standard Benjamini-Hochberg correction. The mass spectrometry proteomics data have been deposited to the ProteomeXchange Consortium via the PRIDE partner repository with the dataset identifier PXD045670 and 10.6019/PXD045670.

## Results

### The *sa43718* allele is embryonic lethal

The sa43718 allele results in a single base pair substitution in exon 6, which is predicted to result in a premature stop codon (Fig 1A). Exon 6 encodes part of the extracellular domain. Based on the sequence change present in the sa43718 allele, we developed a restriction enzyme based genotyping strategy. The sa43718 allele abolishes an XhoI cleavage site and therefore cannot be cleaved by XhoI in a restriction digest. We performed PCR amplification using primers that flank the substituted nucleotide and used restriction digest to identify wildtype, heterozygous, and homozygous larvae. DNA from wildtype siblings was completely digested as predicted (Fig 1B, lane 2), while DNA from homozygous larvae was completely undigested, and DNA from heterozygous carriers was partially digested (Fig 1B, lane 4 and 3, respectively). Homozygous carriers of the sa43718 allele expressed reduced levels of *gabra1* mRNA transcript relative to wildtype siblings (Fig 1C, p<0.05), which was associated with early lethality. No homozygous larvae survived beyond 22 DPF (Fig 1D). These data correspond with mRNA and survival data previously published by Samarut and colleagues using a different nonsense allele ^17^.

**Fig 1.**
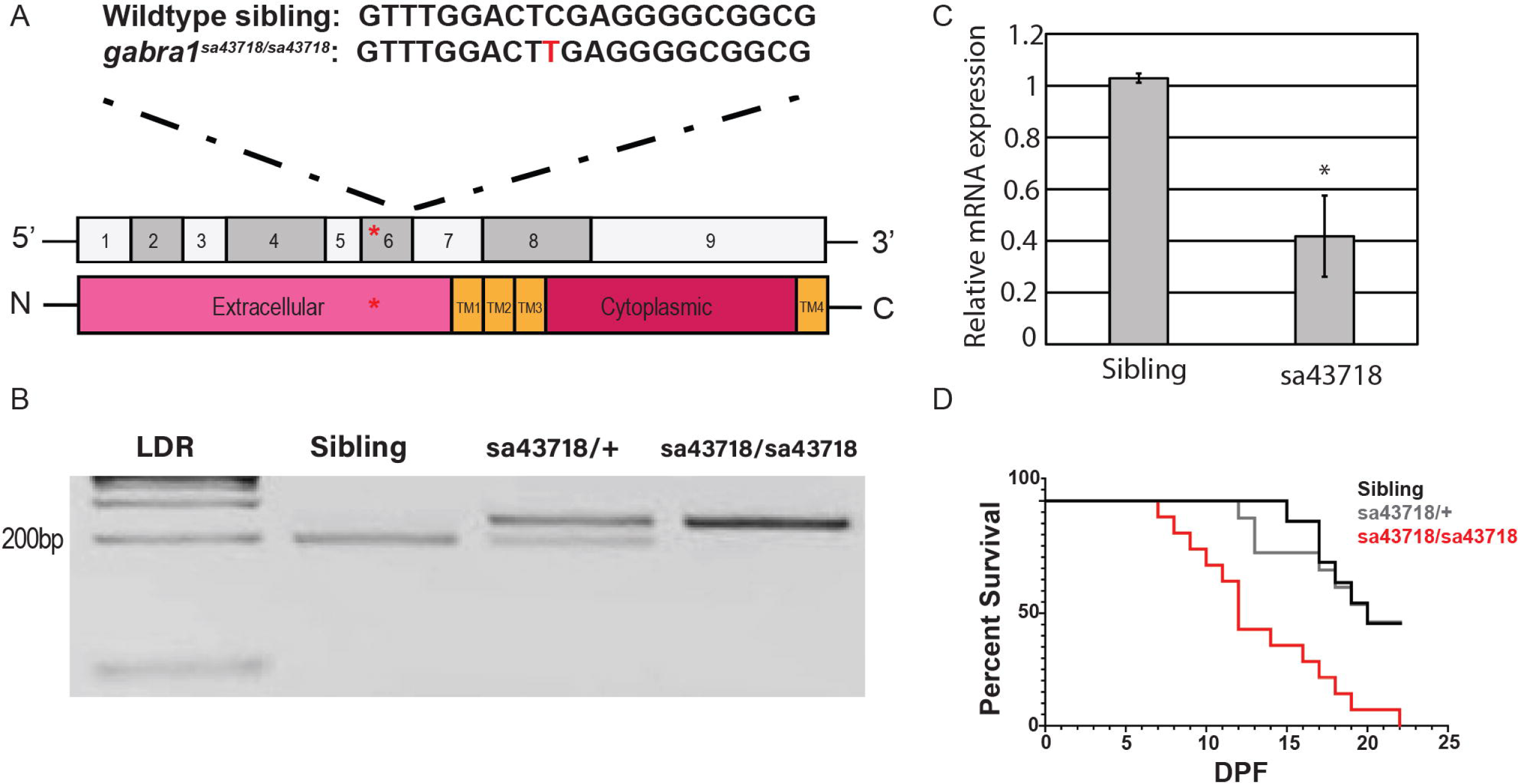
The *sa43718* allele carries a single nonsense mutation in the *gabra1* gene. (A) Schematic representation of the *gabra ^sa^*^43718^ allele. Homozygous individuals of the *sa43718* allele (*gabra1^sa43718/sa43718^*) carry a single base pair substitution (C>T) in the *gabra1* gene (ENSDART0000010000), predicted to result in a premature stop. Schematic of exon structure of the *gabra1* gene, not to scale. The *sa43718* site is indicated with a red asterisk within exon 6, which encodes the extracellular domain of Gabra1. (B) Restriction enzyme digest gel showing digestion of PCR fragments of genomic *gabra1.* The *sa43718* allele abolishes the Xhol recognition site. Therefore, digestion of homozygous (sa43718/sa43718) sample results in an uncut fragment, whereas digestion of wildtype (sibling) sample results in a fully digested fragment. LDR: 1kb plus ladder. (C) Quantitative PCR (qPCR) analysis of the relative expression of *gabra1* in sibling wildtype and homozygous carriers of the *sa43718* allele. Error bars represent the standard error of the mean of biological triplicates. T-test was used to detect statistical significance and *p<0.05. (D) The offspring of *gabra1^sa^*^43718^*^/+^* fish was genotyped by tail-clipping at 3 days post fertilization (DPF). Larvae were raised separately according to their genotypes (sibling n=11, sa43718/+ n=13, sa43718/sa43718 n=14), and their survival was monitored until day 22. Premature death of homozygous carriers of the *sa43718* allele (sa43718/sa43718) was observed starting at day 7 post fertilization, when compared to the wildtype (sibling) and heterozygous (sa43718*/+)* siblings.

### Mutation of *gabra1* results in hypermotility

We have previously reported hypoactivity at 5 DPF upon knockdown of *gabra1* ^2^. In contrast, seizure-like behavior at early and juvenile stages has been reported in a zebrafish model of *gabra1* loss of function ^17^. To begin to rectify these differences, we monitored swimming behavior after a light stimulus, using a similar approach as Samarut and colleagues ^17^. We measured baseline activity for a period of 5 minutes in the dark after a 1-hour acclimation period. During this time, we did not observe any differences in baseline swimming patterns according to total distance swam (mm) or swim speed (mm/s) in the sa43718 allele (Fig 2A&A’). However, after light stimulus, we detected a marked increase in swim speed in the *sa43718* allele (Fig 2A&A’). Swim patterns were increased in mutant larvae during the entire light stimulus (Fig 2C&C’). Importantly, increased speed (velocity) was associated with rapid circling behavior, hyperactivity burst, and whirlpool behavior (Fig 2B) indicative of a seizure-like behavior as defined by the zebrafish behavioral glossary ^19^. Because we did not observe behavioral changes in heterozygous carriers in comparison to wildtype siblings all data shown here are from sibling wildtype or homozygous carriers of the *sa43718* allele. Following the initial burst movement, homozygous larvae have a decrease in speed which normalizes with that of sibling wildtype (Fig S1). Interestingly, because the larvae reduce speed from 660 seconds onward, they respond to a second light stimulus at a similar magnitude of speed as observed in the first light stimulus, with a maximum speed of 10mm/s (900-960 seconds) in each light stimulus (Fig S1).

**Fig 2.**
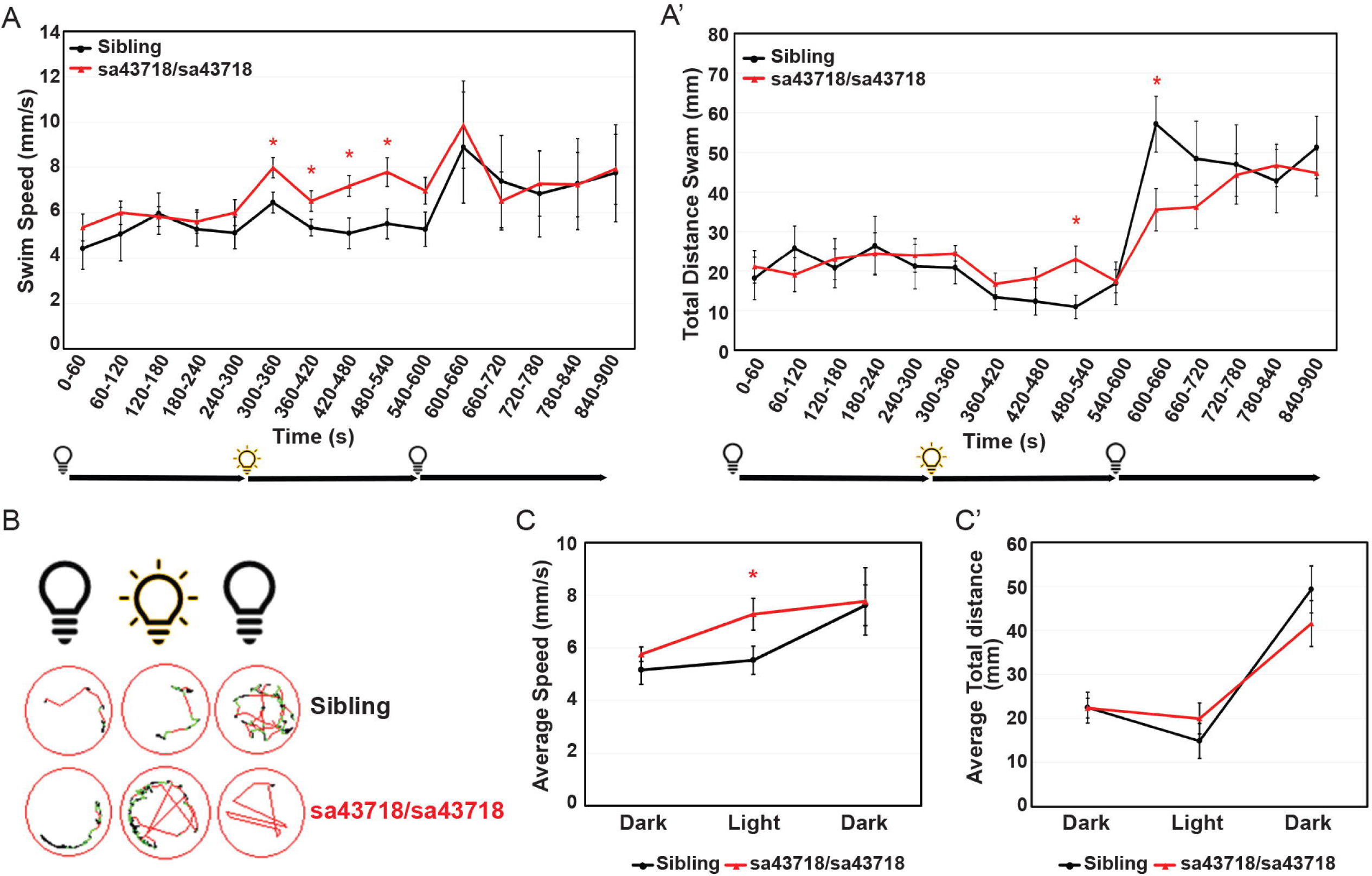
*gabra1^sa43718/sa43718^* larvae undergo seizure-like behavior upon light stimuli. (A & A’) Behavioral analysis of *sa43718* allele and wildtype siblings at 5 days post fertilization. (A) Swim speed (mm/s) over 15 minutes (900 seconds) in dark-light-dark transitions [5 minutes (0-300 seconds) in the dark, 5 minutes (300-600 seconds) in the light, and 5 minutes (600-900 seconds) in the dark) was analyzed using the Zebrabox technology. Data was collected every minute (60 seconds). *gabra1^sa43718/sa43718^* larvae undergo hyperlocomotion upon light stimuli when compared to their wildtype siblings (Sibling). *p<0.05. (A’) Total distance swam over 15 minutes in dark-light-dark transitions. Homozygous carriers (sa43718/sa43718) larvae showed increased distance swam across the light phase, showing a statistically significant increase at seconds 480-540 (minute 9), when compared to their wildtype counterparts (sibling). *gabra1^sa43718/sa43718^* larvae then show decreased distance swam under dark conditions (600-660s) when compared to their wildtype siblings (sibling) *p<0.05. (B) Representative images of swimming track of a single larvae, representing each genotype, generated from the Viewpoint Zebralab Tracking software in dark (180-240s), light (480-540s), and dark (600-660s) conditions. Green lines indicate movements between 4-8mm/s and red lines indicate burst movements (>8mm/s). (C) Average total speed representing overall behavioral responses in dark-light-dark conditions of *gabra1* wildtype (sibling) and homozygous (sa43718/sa43718) larvae. Homozygous carriers (sa43718/sa43718) showed increased average swim speed in light conditions when compared to their wildtype (sibling) counterparts. (C’) Average total distance swam representing overall behavioral responses in dark-light-dark conditions of *gabra1* wildtype (sibling) and homozygous (sa43718/sa43718) larvae. No statistically significant changes were observed in homozygous larvae (sa43718/sa43718) when compared to their wildtype siblings for average distance measurements. All behavioral analysis was performed using a minimum of 30 larvae per biological replicate, over three independent biological replicates. Representative analysis of a single biological replicate is shown here. Sibling (n=13) and *gabra1^sa43718/sa43718^* (n=26).

### Nonsense mutation of Gabra1 does not preclude response to PTZ

We have previously demonstrated that morpholino mediated knockdown of *gabra1* does not prohibit receptor responses to PTZ, a GABA_A_R antagonist ^2^. Therefore, we tested the ability of homozygous carriers of the *sa43718* allele to respond to PTZ. As shown in Fig 3A, treatment with PTZ induces increased swimming distance and speed in wildtype larvae. Within the first five minutes post treatment, in dark conditions, wildtype larvae exhibit an average 2-fold change in total distance swam and a 1.54-fold change in swim speed (Fig 3C and D; gray bars in the dark). The sa43718 allele responded to PTZ but had a blunted response relative to wildtype siblings. For example, the response to PTZ in dark conditions (Fig 3C-D: red bars in dark conditions compared with gray bars). We continued to monitor the speed and distance swam after light stimulus in PTZ treated larvae, both wildtype and homozygous mutants. After light stimulus, wildtype sibling larvae showed a 1.5-fold change in average swim speed and a 2.5-fold change in average total distance swam (Fig 3C-D; light conditions gray bars). In contrast, the sa43718 allele exhibited a 1.4-fold change in average swim speed and 1.7-fold change in average distance swam in response to PTZ treatment, neither was statistically significant (Fig 3C-D). The reduced response to PTZ treatment by the sa43718 allele was statistically significant in dark conditions for speed (p=0.03) and distance (p=0.002) indicating a blunted response.

**Fig 3.**
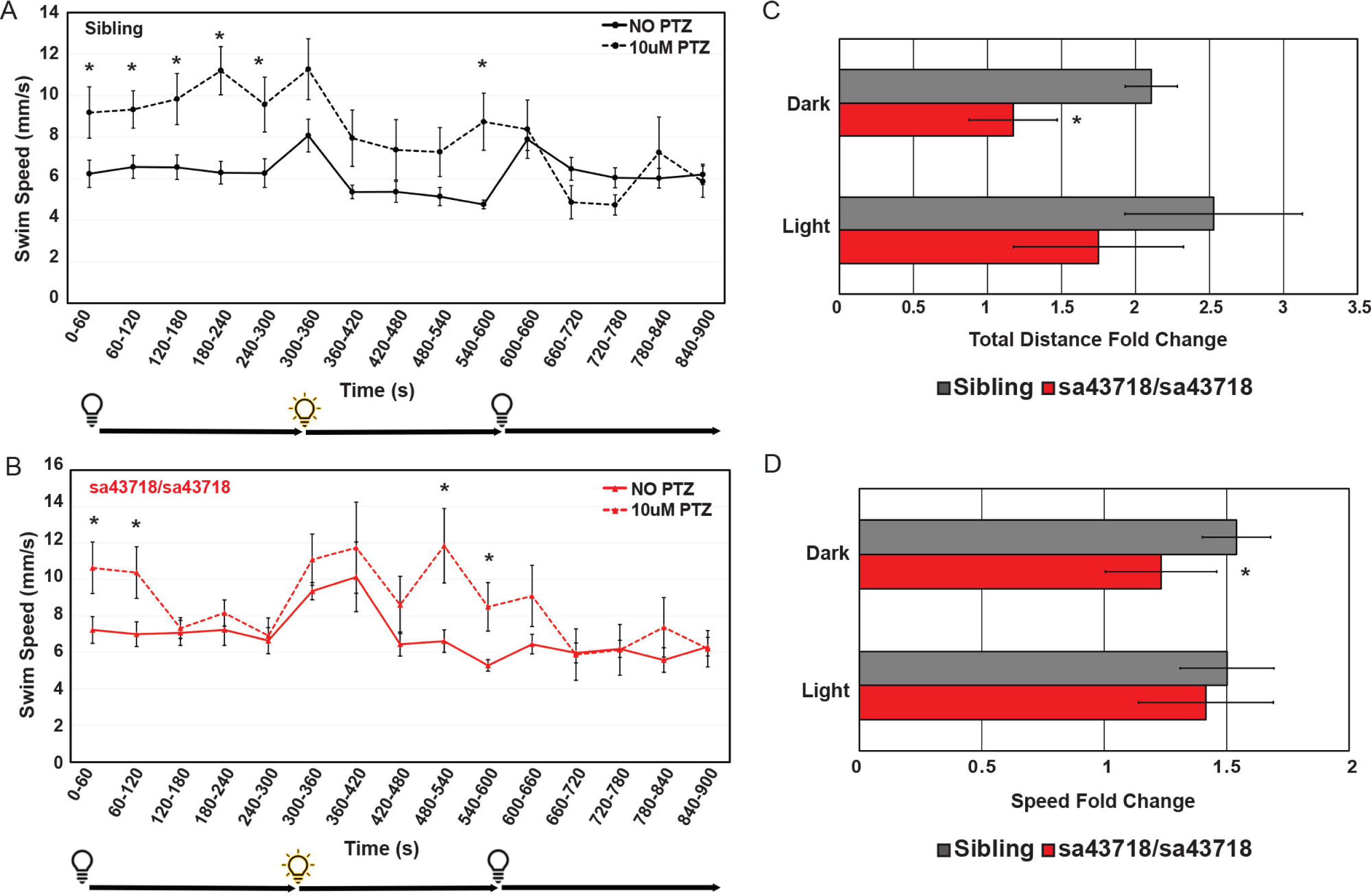
Locomotion response to pentylenetetrazole (PTZ) in the *sa43718* allele. (A-B). Behavioral responses of the *sa43718* allele at 5 days post fertilization (DPF) were analyzed using the Zebrabox. (A) Comparison of wildtype sibling (Sibling) larvae response to treatment with 10uM PTZ (10uM PTZ) or no treatment (NO PTZ) at 5 days post fertilization. NO PTZ n=18, 10uM PTZ n=16. *p<0.05 (B) Comparison of homozygous carriers (sa43718/sa43718) response to treatment with 10uM PTZ (10uM PTZ) or no treatment (NO PTZ) at 5 days post fertilization. NO PTZ n=13, 10uM PTZ n=14. *p<0.05. Swim speed (mm/s) was analyzed using the distance swam and swimming duration data generated by ViewPoint software. Larvae were monitored for 5 minutes in the dark (0-300 s), 5 minutes in the light (300-600 s), and 5 minutes in the dark (600-900 s). Data was collected every minute (60 s). Error bars represent the standard error of the mean. (C) Average fold change (total distance) response to treatment with PTZ in dark (0-300s) and light (300-600s) conditions by wildtype (sibling) and sa43718/sa43718 mutants. *p=0.002. (D) Average fold change (swim speed) response to treatment with PTZ in dark (0-300s) and light (300-600s) conditions by wildtype (sibling) and sa43718/sa43718 mutants. *p=0.03. Error bars represent standard deviation in C-D.

### Expression of GABA_A_ receptor subunits in the sa43718 allele

Previous studies suggest that mutation of *Gabra1* in mice ^13^ and zebrafish ^2, 17^ results in abnormal expression of other subunits of the GABA_A_R. Thus, we hypothesized that the expression of other GABA_A_R receptor subunits were disrupted in the *sa43718* allele. We first measured the expression of β2 and γ2 subunits, as these are present in the most common form of the GABA_A_R receptor, alongside α1. We did not observe abnormal expression of the transcripts encoding either the β2 or γ2 subunits (Fig 4A). Consequently, we hypothesized that a version of the GABA_A_R receptor is being produced. We next measured the expression levels of α2-6. We observed a statistically significant decrease in α2, α3, and α5, however, the decrease was not more than 2-fold in any case (Figure 4A).

**Fig 4.**
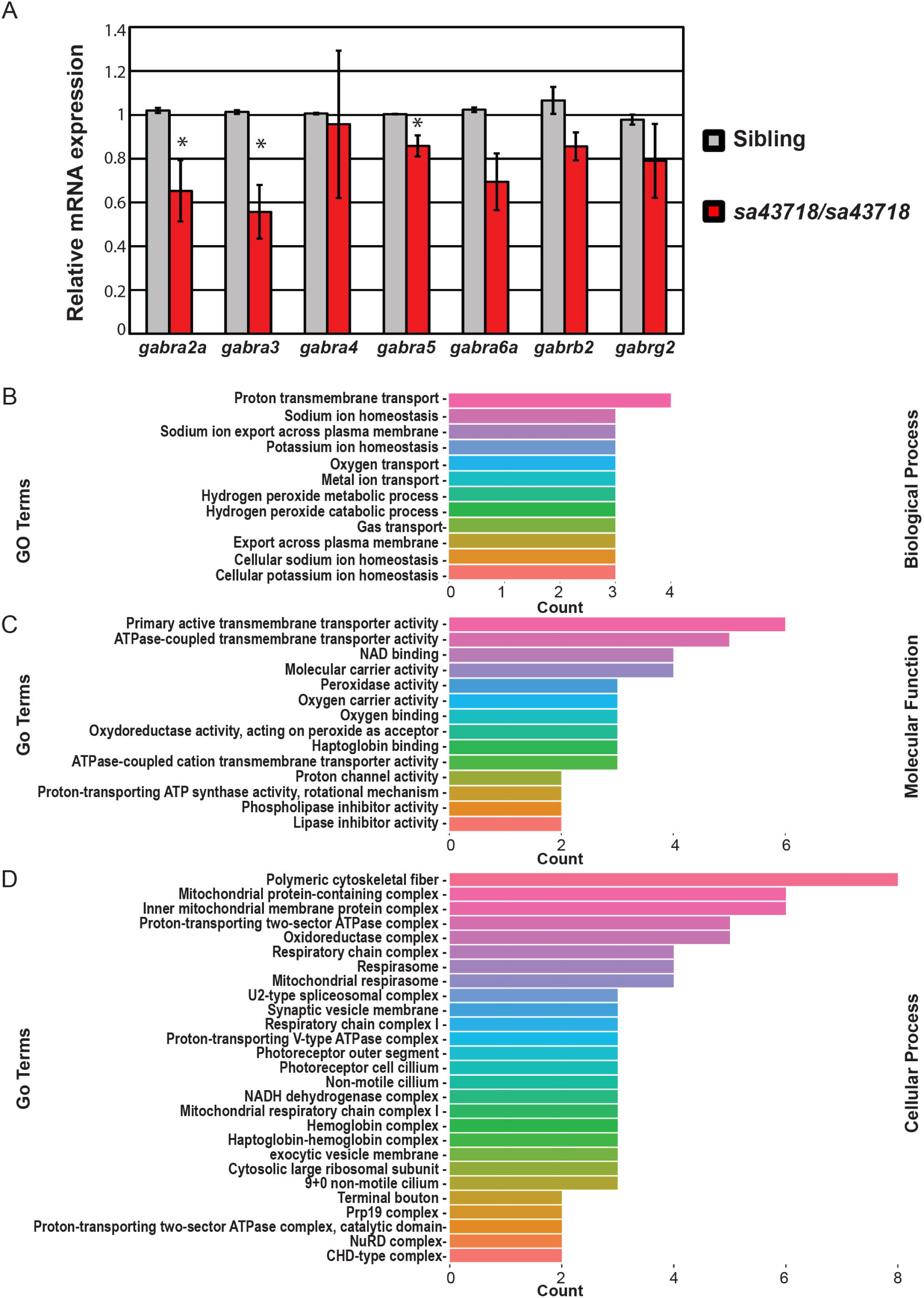
Differential Gene and Protein Expression of *sa43718* allele carriers. (A) Relative expression of major subunits of the GABA_A_R. *gabra2a* (α2), *gabra3* (α3), *gabra4* (α4), *gabra5* (α5), *gabra6a* (α6a), *gabrb2* (β2), and *gabrg2* (γ2) were analyzed in wildtype siblings (sibling) and homozygous (sa43718/sa43718) mutants. Graph represents the mean relative mRNA expression obtained from 3 biological replicates each with a pool of N=10-12 larvae per group. * p<0.05 Errors bars represent standard error of the mean from biological replicates. (B-D) Gene Ontology terms enriched by proteins with significant expression (p-value <0.05) between wildtype sibling and *sa43718* homozygous mutant larvae at 5 days post fertilization. Terms are separated by (B) Biological Process, (C) Molecular Function, (D) Cellular Process.

### Proteomic analysis identifies differentially expressed proteins (DEPs) in the *sa43718* allele

We uncovered normal expression of β2 and γ2 transcript levels with mild to moderate changes in the expression of other α subunits. We next hypothesized that these changes in expression are associated with abnormal inhibitory network development, which was previously described. To begin to investigate this hypothesis, we isolated protein from whole brain homogenates and performed proteomics analysis. A total of 3500 proteins were identified in the wildtype sibling brain homogenate, 3511 were identified in heterozygous animals, and 3552 were identified in the homozygous carriers of the sa43718 allele. A complete list of identified proteins is in S1 file. From these proteins, we identified a total of 173 DEPs in homozygous mutants relative to wildtype clutch mates. Of these 81 were up-regulated and 93 were down-regulated. A complete list of DEPs can be found in file S2. DEPs of interest include Synaptotagmin, Complexin 4a, Dynactin subunit 1, Dynamin GTPase, Secretogranin II, and a Vat1 homolog. Each of these proteins has a function in synaptic fusion, transport, exocytosis, endocytosis, or direct interactions with small synaptic vesicles. Interestingly, we also noted a significant number of DEPs involved in microtubule growth, microtubule binding, vesicle recycling, retrograde transport along microtubules, or microtubule folding. These include microtubule associated protein, RP/EB family member 3b, Dynactin subunit 1, Dynamin GTPase, and Tubulin-specific chaperone A. We further annotated DEPs by biological process (Fig 4B), molecular function (Fig 4C), and cellular process (Fig 4D). According to biological process, DEPs were clustered in Gene Ontology (GO) groups associated with proton transport including sodium and potassium export across the plasma membrane. Some examples of DEPs in this category include Slc82b, Atp1a1b, Atp1b3a, and Atp1a1a.3. All these DEPs are important for sodium and potassium homeostasis consistent with GO analysis. GO analysis by molecular function also indicated molecular carrier activity and active transport mechanisms, consistent with the GO analysis according to biological process. GO analysis by cellular process indicated DEPs in the respiratory chain, mitochondrial protein containing complexes, the cytoskeleton, proton transport, and exocytosis. These data collectively indicate mitochondrial transport, synaptic vesicle regulation, and proton homeostasis are dysregulated after mutation of Gabra1.

## Discussion

The GABA_A_R has previously been associated with seizure and epileptic phenotypes ^1, 9, 16^. In support of this notion, we previously reported a heterozygous missense mutation in humans that is associated with a seizure phenotypes ^2^, which has subsequently been shown to be a loss of function allele ^12^. Multiple model systems have been developed to study early *GABRA1* function. These include mouse and zebrafish models. Survival of homozygous knockouts in mice and zebrafish is compromised ^13, 17^. Due to external fertilization and early juvenile survival of homozygous mutants, zebrafish provide a window of opportunity to analyze developmental and juvenile onset of seizures ^20–25^. However, in previous studies germline nonsense mutation of zebrafish *gabra1* and morpholino mediated knockdown of *gabra1* demonstrated different behavioral phenotypes after a light stimulus ^2, 17^. Here we characterized the phenotypes of a novel allele, known as the *sa43718* allele. The *sa43718* allele is a single base pair change in exon 6, which results in a premature stop codon and leads to a decrease in *gabra1* expression. The *sa43718* allele reduces viability and homozygous mutants die by 22 DPF. This is consistent with the viability of other zebrafish ^17^ and mouse models ^14^.

We observed seizure like (hypermotility, whirlpool behavior) behavior upon light stimulus in the *sa43718* allele. These data are supported by previously published works, which established seizure like behavior at the juvenile stage in a zebrafish germline nonsense mutant ^17^. Similar results have also been documented in mouse models of *Gabra1* mutation, although these data are sex and strain specific ^13–15^. In our experiments, the zebrafish larvae have not yet determined sex and therefore, sex is not a variable considered in our experiments. Hyperactivity and seizure like behavior in the *sa43718* allele contrasts with our previous study using morpholino mediated knockdown ^2^ but validates findings from others using another nonsense allele. Morpholinos are known to have off-target effects ^26^, however our previous study found hypomotility using two independent morpholinos. It is a strong possibility that dosage, timing, and level of knockdown could play a role in the interpretation of these results. Nonetheless, our analysis of *sa43718* provides strong evidence, particularly when collectively considered with data from Samarut and colleagues, that nonsense mutation of *gabra1* is lethal and is associated with the early onset of seizure like behavior.

Previous studies also demonstrated that the overall brain morphology of *gabra1* mutant brains was normal. This is consistent with functional analysis of the p.T292I patient variant we discovered. Chen and colleagues established that the pT292I variant can produce a receptor in HEK293 cells, but reduces channel open time ^12^. Interestingly and despite a normal morphology, Samarut and colleagues showed 460 differentially expressed genes in *gabra1* mutant brains. Of these genes, 3 GABA_A_R subunits, in addition to *gabra1*, were down regulated ^17^. Consistent with these data, we observed reduced expression of other α subunits, specifically α2, α3, and α5. However, none of the α subunits were reduced more than 50%, suggesting that some transcripts are produced and could be incorporated into the GABA_A_R. Incorporation of these subunits, albeit at a lower level, is likely underlying the ability to respond to PTZ that we observed in homozygous mutants.

Though the *sa43718* allele responded to PTZ the overall response was blunted relative to wildtype siblings and the transcript levels of other α subunits was mildly reduced so we hypothesized the loss of *gabra1* caused abnormal development of the inhibitory system. Therefore, we performed proteomics analysis to determine if the behavioral effects are associated with differential protein expression of proteins required for synapse development. We did not identify a single GABA_A_R subunit differentially expressed in the *sa43718* allele at the protein level. This is attributed to low overall protein expression in the brain homogenate and detection. Therefore, we focused our analysis on proteins that were abundantly and consistently detected in technical (3) and biological replicates (2). We identified proteins with functions in synaptic vesicle transport, microtubule organization/growth, mitochondrial function, and regulation of potassium and sodium homeostasis as differentially expressed. These biological categories are consistent with data from Samarut and colleagues, who noted abnormal mRNA expression of genes in neuronal function and mitochondrial function, among others ^17^. However, we did not detect a high percentage of overlapping genes and proteins between our two studies, though biological processes did overlap, with significant emphasis on synaptic vesicles, synaptic function, and the mitochondria. Of note, we identified abnormal expression of synaptotagmin, secretogranin II, and synapsin IIb; all of which are important for synaptic vesicle release at the presynaptic membrane. Synapsin I was also downregulated and has been associated with epileptic phenotypes ^27–30^. Furthermore, we identified downregulation of proteins implicated in mitochondrial complex I deficiency, including NADH:ubiquinone oxidoreductase core subunit S2 and NADH:ubiquinone oxidoreductase core subunit A10. Remarkably, mutations in the genes encoding these proteins are associated with Leigh syndrome, one of the most common mitochondrial diseases, in which epilepsy is a primary phenotype ^31, 32^. Interestingly, mutation of *GABRA1* in humans was prematurely diagnosed as a primary mitochondrial disorder prior to exome sequencing ^24^.

Here we have characterized an additional germline mutant of the zebrafish *gabra1* gene. We observed early lethality, seizure-like hypermotility after light stimuli, and abnormal expression of proteins involved in vesicle transport, synaptic vesicle release, exocytosis, and endocytosis. Our data support the hypermotility and abnormal development of inhibitory synapses described previously in a zebrafish nonsense mutant ^17^. Collectively, these data suggest that models derived from germline mutation of *gabra1* rather than transient knockdown are more physiologically relevant.

### Limitations of the study

The present study characterizes the *sa43718* allele. However, due to the lack of antibodies, we were unable to detect Gabra1 protein in zebrafish and we could not validate the decrease of Gabra1 using proteomics due to low detection levels. None of the subunits were consistently detected by proteomics or western blot. Thus our study relies on mRNA validation of *gabra1* expression and sequencing to confirm the mutation of interest. While the mutation can be confirmed using Sanger sequencing there still exists the potential for off-target effects. However, our analysis recapitulates and validates previous studies in zebrafish with a germline mutant of *gabra1*.

## Acknowledgements

The authors would like to thank the members of the Quintana Lab, past and present, for discussions, animal care, genotyping, and general laboratory maintenance. A special thank you to Pamela Rodriguez, Valeria Virrueta, and Carla Canales for their efforts in the laboratory and their explicit help with genotyping embryos related to this manuscript. The work provided herein could not be completed without the support of The University of Texas El Paso core facilities and Border Biomedical Research Center (BBRC).

## Supporting information

**S1 File. Complete list of identified proteins.**

**S2 File. Complete list of differentially expressed proteins.**

**S3 File. Supplemental figure and figure legend.**

## Author contributions

DP analyzed data, performed RNA analysis, qPCR, protein analysis (western blots not included due to low cross reactivity of antibodies), confirmatory and validatory experiments. NGR performed experiments, provided experimental conceptualization, wrote aspects of the manuscript, performed zebrafish care and maintenance, supervised trainees that provided input/data analysis or acquisition. AMQ performed data analysis, wrote portions of the manuscript, obtained funding, provided mentorship to NGR, and performed data analysis. BEP performed validatory analysis of behavior phenotypes. IP performed survival analysis and produced graphs related to survival. BG, ILE, and CCE performed mass spectrometry, protein isolation, and data analysis for proteomics. CG and AG performed critical genotyping and analysis to complete the manuscript and VLC provided optimized protocol for protein isolation and data analysis.

## Data availability

All data and reagents are available upon request from the corresponding author.

