## Supplementary material for "Characterization of the zebrafish *gabra1^sa43718/sa43718^* germline loss of function allele confirms a function for Gabra1 in motility and nervous system development": S3 Supplemental Figures

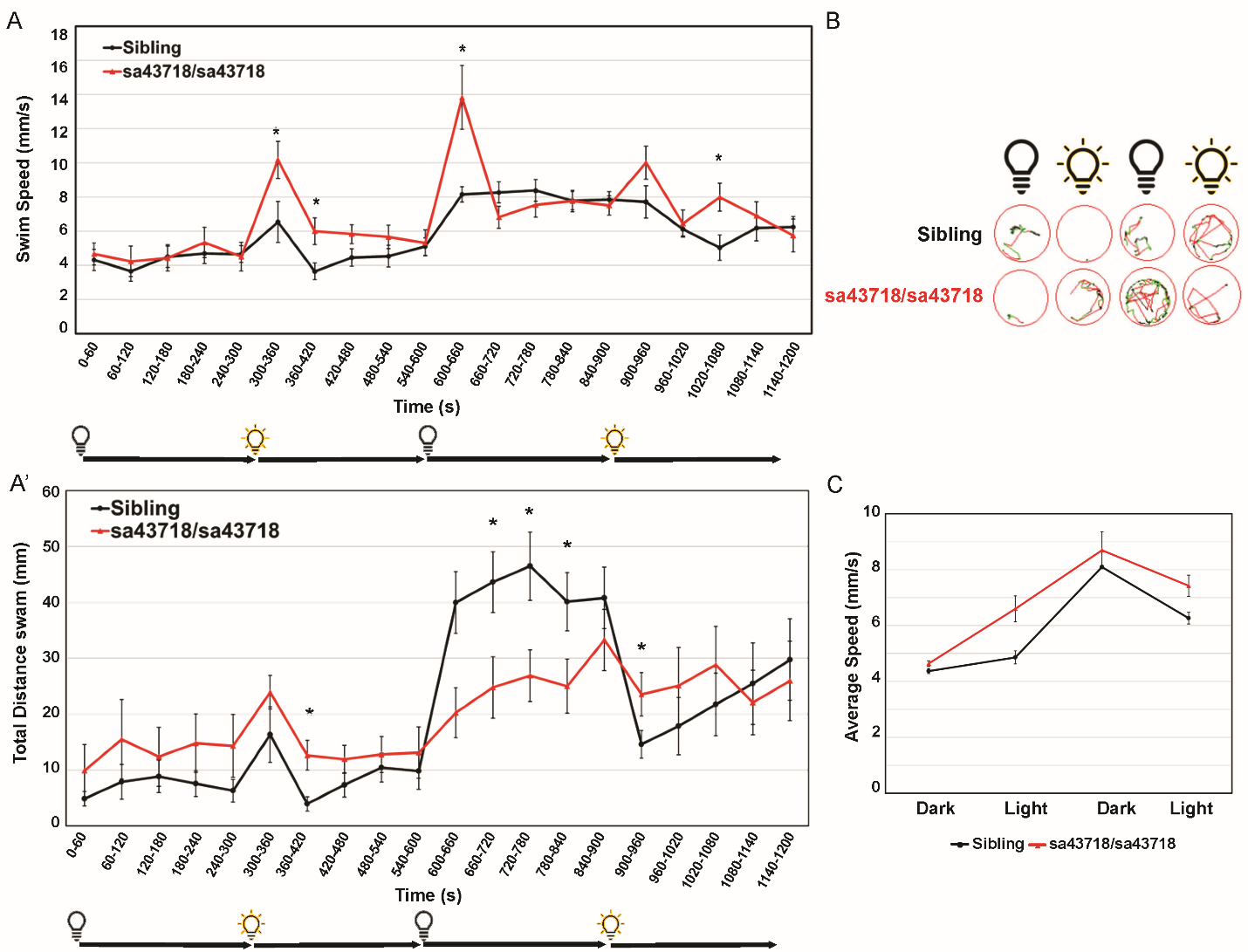

**S1 Figure. Light stimuli induce hyperactive locomotion of *gabra1^sa43718/sa43718^* larvae.** A & A’. Behavioral analysis of gabra1*^sa43718/+^* offspring at 5 days post fertilization. A. Swim speed (mm/s) over 20 minutes (1200 seconds) in dark-light-dark-light transitions [5 minutes (0-300 seconds) in the dark, 5 minutes (300-600 seconds) in the light, 5 minutes (600-900 seconds) in the dark, and 5 minutes (900-1200 seconds) in the light] by 5-day post fertilization (DPF) larvae, analyzed using the Zebrabox technology. Data was collected every minute (60 seconds). *gabra1^sa43718/sa43718^* larvae exhibit hyperlocomotion upon light stimuli when compared to wildtype siblings (Sibling). *p<0.05. A’. Total distance swam over 20 minutes in dark-light-dark-light transitions. Homozygous carriers (*sa43718/sa43718*) larvae showed increased distance swam across the light periods (seconds 300-600 and 900-1080), when compared to wildtype siblings (sibling). Homozygous larvae then show decreased distance swam under dark conditions (seconds 600-900) when compared to wildtype siblings (sibling) *p<0.05. B. Representative images of swimming track of a single larvae, representing each genotype, generated from the Viewpoint Zebralab Tracking software in dark (0-60s), light (300-360s), dark(600-660s), and light (1020-1080s) conditions. Green lines indicate movements between 4-8mm/s and red lines indicate burst movements (>8mm/s). C. Average total speed representing overall behavioral responses in dark-light-dark conditions of *gabra1* wildtype (sibling) and homozygous (sa43718/sa43718) larvae. Homozygous carriers (sa43718/sa43718) showed global increased average swim speed when compared to wildtype (sibling) counterparts. Error bars represent the standard error of the mean. *Wildtype sibling (n=21) and gabra1^sa43718/sa43718^ (n=19).*
